# A Novel Pathway Implicated in Regulating Cognitive disfunction in a *Drosophila* Alzheimer’s Disease Model through Acer Inhibition and CG2233 Modulation

**DOI:** 10.1101/2023.12.21.572837

**Authors:** Judy Ghalayini, Shin-Hann Lee, Oxana Gluscencova, Konstantin G. Iliadi, Gabrielle L. Boulianne

**Affiliations:** Program in Developmental & Stem Cell Biology, Peter Gilgin Center for Research & Learning, The Hospital for Sick Children, Toronto, Ontario, Canada, M5G 0A4; Department of Molecular Genetics, University of Toronto, M5S 1A1

## Abstract

Alzheimer’s disease (AD) is a progressive neurodegenerative disorder, accounting for most dementia cases worldwide. Current therapies for AD have limited effectiveness in slowing disease progression or delivering a cure. As such, there is an immediate need for ongoing research and innovative strategies to tackle this multifaceted disease. Recently, several studies have implicated the renin-angiotensin system (RAS), known to regulate blood pressure, as a possible therapeutic target for AD. RAS-inhibiting drugs, including angiotensin-converting enzyme inhibitors (ACE-Is), have been shown to reduce the incidence and progression of AD. However, the literature describing their beneficial effects is inconsistent, with contradictory findings reporting no effects. How these drugs may function in AD remains poorly understood. Our previous work in *Drosophila* models expressing AD-related transgenes investigated the benefits of captopril, an ACE-I, and found it effectively rescued AD-related phenotypes including cognitive performance independent of Aβ42 changes. Importantly, our study implicated Acer, a homolog of mammalian ACE, as a key player. In our current study, we demonstrate that the beneficial outcomes of Acer inhibition depend on preventing its catalytic activity and downstream target processing. We identify CG2233 as a prospective target and reveal its functional interaction with Acer. Furthermore, we show CG2233 is implicated in AD-related pathways in Aβ42 expressing flies. Together, these findings provide a new avenue to study the role of ACE in AD.

**Significance Statement:** AD is a devastating neurodegenerative disorder with limited therapeutic success. Emerging research highlights the potential of inhibiting the renin-angiotensin system (RAS) in AD. Epidemiological findings and experimental studies have shown promising outcomes with RAS-targeting drugs including angiotensin-converting enzyme inhibitors (ACE-Is). Our previous work in *Drosophila* AD models revealed the efficacy of captopril, an ACE-I, in improving AD-related phenotypes. Moreover, we identified Acer as a key player in these mechanisms. Our current study further elucidates the role of Acer, identifies CG2233 as a potential target, and uncovers their functional interaction, shedding light on pathways relevant to AD phenotypes. This research underscores the significance of investigating ACE and ACE-I mechanisms in AD, offering potential innovative means for AD therapy.

## Introduction

Alzheimer’s disease (AD) is a progressive neurodegenerative disorder accounting for the majority of dementia cases worldwide (Barker et al., 2002). It is characterized by two pathological hallmarks, amyloid plaques and neurofibrillary tangles, composed of aggregates of extracellular β-amyloid (Aβ) peptide and intracellular hyperphosphorylated tau proteins, respectively (Holtzman et al., 2011). Among various proposed mechanisms for AD pathogenesis, the amyloid cascade has been a leading hypothesis suggesting that pathological accumulation of longer, more toxic forms of Aβ peptide, Aβ42, which are associated with brain atrophy, regional hypometabolism, synaptic dysfunction, neuro-inflammation and oxidative stress initiates and drives AD (Hardy & Selkoe, 2002). This hypothesis has driven drug development efforts for over two decades with limited success (Huang et al., 2020). Only recently have drugs designed to target and clear Aβ been approved as treatments for AD, although with limited impact on cognitive improvement (Reardon, 2023; Tampi et al., 2021). Consequently, there is ongoing need to develop therapies that prioritize targets other than Aβ.

A growing body of evidence suggests that pharmacological inhibition of the renin-angiotensin system (RAS), a critical regulator of blood pressure and fluid balance, may have a positive impact on AD. Early epidemiological studies led the investigation demonstrating a beneficial connection between RAS-targeting anti-hypertensive medications, such as angiotensin-converting enzyme inhibitors (ACE-Is) and angiotensin receptor blockers (ARBs), and AD (Barthold et al., 2018; Davies et al., 2011; Deng et al., 2022; Hajjar et al., 2005; Hajjar et al., 2012; Li et al., 2010; Ouk et al., 2021). Notably, these studies revealed that patients treated with ACE-Is or ARBs exhibited a lower incidence and slower progression of AD compared to those using alternative anti-hypertensive drugs, although the mechanism of action remains unclear. Furthermore, these beneficial outcomes have been supported in experimental studies conducted in mouse models of AD. Several studies have shown that ACE-Is and ARBs lead to improvements in cognitive function in AD mouse models, accompanied by reductions in AD-related pathological features such as neuroinflammation, oxidative stress and Aβ deposition (AbdAlla et al., 2013; Asraf et al., 2018; Dong et al., 2011; Ongali et al., 2014; Royea et al., 2017; Torika et al., 2016). However, it is worth noting that some research contradicts these findings, with studies suggesting that ACE may cleave Aβ peptides and others demonstrating an increase in Aβ deposition following treatment with ACE-Is (Liu et al., 2019; Zou et al., 2007). Consequently, understanding the protective role of ACE-Is in AD requires further investigation and unravelling their mechanisms of action is important in accomplishing this goal.

In our previous work, we investigated the impact of captopril, an ACE-I, in *Drosophila* models of AD. We showed that captopril suppressed a rough eye phenotype and reduced cell death in the brains of flies expressing AD-related transgenes, including Aβ42 (Lee et al., 2020). Additionally, we demonstrated that captopril ameliorated memory deficits observed in these flies. Similar findings were also reported by (Thomas et al., 2021). Importantly, our observations indicated that captopril had no discernible effect on the levels or clearance of Aβ42, nor did it influence degenerative phenotypes in *Drosophila* models expressing human tau. Furthermore, our research demonstrated that a null mutation in *acer*, a homolog of mammalian ACE, recapitulated the beneficial outcomes found with captopril, suggesting that Acer is its target (Lee et al., 2020). Noteworthy, downstream RAS substrates of ACE are not found in *Drosophila* (Fournier et al., 2012), suggesting that Acer functions through a novel target.

Here, we demonstrate that the beneficial outcomes of Acer inhibition observed in Aβ42 expressing flies is dependent on he prevention of its catalytic activity, potentially underscoring the significance of interfering with the processing of its downstream target. Furthermore, we identified CG2233 as a putative target and demonstrated a functional interaction between the two proteins. Finally, we show that CG2233 is involved in pathways that contribute to AD-related phenotypes in Aβ42 expressing flies. Altogether, our findings suggest a new pathway that can be dissected further in *Drosophila* to provide insight into the role of ACE and ACE-Is in AD.

## Materials and Methods

### *Drosophila* stocks and husbandry

All fly stocks were reared on standard medium (cornmeal, yeast, agar, and molasses) at 25°C and 60%–65% humidity on a 12-hour light/12-hour dark cycle. Fly strains obtained from the Bloomington *Drosophila* Stock Center include *Canton-S* (CS), *w^1118^*, *elav-Gal4^C155^* (#458), *UAS-APPAbeta42.B* (#33769) (referred to as *UAS-Aβ42*), vas-Cas9(III) (#51324) and Tub-PBac (#8283). *Acer^Δ168^* allele (*acer^null^*) flies were gifted by Dr. A. Carhan (Carhan et al., 2011)). Fly lines *vas-Cas9*(III) and *Tub-PBac* were used to generate an *acer ^H375K, H379K^* mutant allele (*acer^PD^*) and a *CG2233Δ* mutant allele. Transgenic flies, expressing human *Aβ42* (*UAS-Aβ42)* in neurons driven by *elav-Gal4^C155^* were used as an model of AD (Iijima et al., 2004). To study the effects of an *acer^null^* allele or an *acer^PD^* mutant allele in the background of flies expressing Aβ42, mutations were recombined with *UAS-Aβ42* on the second chromosome and crossed to generate driver lines *elav-Gal4^C155^*; *acer^null^*; + and *elav-Gal4^C155^*; *acer^PD^*; +, respectively. To study the effects of a *CG2233Δ* null mutant allele in the background of flies expressing Aβ42, the mutant allele was recombined with *elav-Gal4^C155^* on the X chromosome and female flies were crossed to *UAS-Aβ42* transgenic males or, *w^1118^* and used as controls, respectively. In all experiments, respective driver lines were crossed to *w^1118^*and served as appropriate controls.

### Generating an *acer^H375K, H379K^* (*acer^PD^*) mutant allele using CRISPR-Cas9 scarless gene editing

The *acer^PD^* mutant allele resulting in substitutions: H375K and H379K was generated using CRISPR/Cas9-mediated homology directed repair (HDR) editing, employing a modified “scarless” strategy (https://flycrispr.org/) (Bier et al., 2018). Two sgRNAs (sgRNA1: TTTACCAAGACAGCGATGTA, sgRNA2: AACAACAGCCGGCTGTCTAC) flanking the targeted genomic sequence were designed using the FlyCRISPR algorithm (http://targetfinder.flycrispr.neuro.brown.edu/) (Gratz et al., 2014) and their efficiency was evaluated using (https://www.flyrnai.org/evaluateCrispr/). Both sgRNAs were cloned into the pCFD4d plasmid (gifted by Dr. T. Hurd, Addgene #83954) using Gibson Assembly with complementary oligos corresponding to each individual sgRNA. The HDR donor plasmid was constructed by incorporating a designed HDR template into the backbone of pScarlessHD-DsRed-w+ plasmid (Addgene #80801). The HDR template comprised the following: 1) ∼1kB 5’ homology arm, upstream of 5’ most sgRNA site, 2) repair template with mutations of interest (C2020A, T2022A, C2032A and C2034A) and single base pair silent mutations (G1971C and G2079T) introduced at PAM sequence sites to prevent Cas9 from cutting the repair template or repaired genome, 3) a 3xP3 DsRed marker cassette flanked by PBac transposon ends and, 4) ∼1kB 3’ homology arm, downstream of 3’ most sgRNA site. PBac transposons target genomic TTAA sites for insertion. The TTAA site is duplicated upon transposon insertion and restored to a single TTAA site upon excision. To achieve a “scarless” excision of the dsRed marker cassette, a single base pair silent mutation (G2154A), which creates a TTAA site upstream of the cassette, was introduced into the genome via the repair template. The HDR template DNA sequence was designed with SnapGene® software (from Dotmatics; available at snapgene.com) using NBCI reference sequence (NT_033779.5) and ordered from Bio Basic. The template was then cloned via Gibson Assembly into the backbone of a pScarlessHD-DsRed-w+ plasmid that was linearized with restriction enzymes SrfI and EcoRI.

The pCFD4d plasmid and HDR donor plasmid were injected into *vas-Cas9* flies by BestGene Inc. The injected flies were balanced and the progeny were screened for DsRed expression in fly eyes. DsRed+ flies were maintained as a stock and sequenced to determine the presence of mutations of interest. DsRed+ fly lines with successful HDR were then crossed to *Tub-PBac* flies, a strain expressing PBac transposase, to excise the DsRed marker cassette leaving behind only the intended mutations of interest. Progeny from the cross were screened and DsRed-flies were selected and established as a stock following verification of scarless removal by PCR sequencing.

### Generating *CG2233Δ* mutant lines

The *CG2233Δ* null mutant allele was generated using CRISPR/Cas9-mediated non-homologous end join (NHEJ). Two sgRNAs (sgRNA1: GGTTACTTCCGTTATGGCCC and sgRNA2: GGACACTTGGTTGCACATCG) flanking the targeted genomic sequence were designed using the FlyCRISPR algorithm (http://targetfinder.flycrispr.neuro.brown.edu/) (Gratz et al., 2014) and their efficiency was evaluated using (https://www.flyrnai.org/evaluateCrispr/). Both sgRNAs were cloned into the pCFD4d plasmid (gifted by Dr T. Hurd, Addgene #83954) using Gibson assembly with complementary oligos that correspond to each individual sgRNA. The pCFD4d plasmid was injected into *vas-Cas9* flies by BestGene Inc. Flies carrying potentially mutated chromosomes were captured over a balancer chromosome and made into stocks. The stocks were subsequently screened for the desired deletion by PCR with a set of deletion-flanking primers, following a previously described screening workflow (https://flycrispr.org/protocols/crrna/). Sequencing of PCR products from generated mutant lines revealed the expected deletion.

### CG2233 polyclonal antibody generation

Custom rabbit polyclonal antibodies recognising CG2233 were generated using PolyExpressTM Premium antigen-specific affinity purified pAb package provided by GenScript. Antibodies were generated against recombinant CG2233 that lacks its N-terminal signal sequence (MFSINAVILGILVTSVMA) by immunizing rabbits with purified 6X His-tagged CG2233 proteins.

### Western blotting

To measure Acer and CG2233 protein levels, whole-flies or adult heads were homogenized in RIPA lysis buffer with protease inhibitor (Roche) using a mortar and pestle. Extracted protein lysates were run on a 10% SDS-PAGE gel and transferred onto a PVDF membrane. Membranes were blocked for 1 hour at room temperature (RT) in 5% milk solution (prepared in Tris-buffer saline with 0.2% Tween-20 (TBST)). Blocking was followed by primary antibody incubation overnight at 4°C with antibodies: rabbit anti-Acer (1:1000) gifted by Dr. A. Carhan (Carhan et al., 2011), rabbit anti-CG2233 (1:1000) and mouse anti-alpha tubulin (1:1000) (abcam, Cat #ab7291). Primary antibody solution was removed, membranes were washed 3x10 min with TBST and incubated with HRP conjugated secondary antibodies: goat anti-rabbit IgG (1:10,000) (Invitrogen, Cat #31460) and goat anti-mouse IgG (1:10,000) (Invitrogen, Cat #31430) for 2 hours at RT. Membranes were washed and developed with ECL (Bio-Rad, Cat #170506) and blots were imaged with Odyssey FC1 (Licor) using Image Studio^TM^ Lite software.

### TUNEL labelling

Brains from aged flies were dissected in cold 0.5% PBS-T (0.5% TritonX-100 prepared with 1xPBS) and fixed in 4% PFA at RT for 30 minutes. Brains were then washed 3x10 min in 0.5% PBS-T at RT and 1x15 min in 0.1% sodium citrate solution at 4°C, followed by 2x10 min washes in 0.5% PBS-T. Brains were then incubated with TUNEL reagent following manufacturer instructions (Roche, in situ cell death detection kit, cat# 11684795910). Following incubation, samples were washed in 1xPBS and mounted using Vectashield. Z-stack images of whole brains were acquired using a Nikon A1R confocal microscope, z-stacks were merged and cell death counts were manually measured.

### Courtship conditioning assay

Experimental male flies were collected within 6 hours after eclosion and individually housed in culture vials containing standard media until the experimental day. 5-day-old *CS* virgin females were mated one day prior to the experiment day and used for both training and testing. Male courtship behavior was observed in custom-made Perspex chambers (15 mm in diameter and 5 mm in height) with a sliding partition that divided the chamber in two halves with respective entries. Chambers were cleaned with 70% ethanol and dried prior to each measurement. To video-record courtship behavior, a single experimental male was placed into a chamber with a mated *CS* female and kept separate for at least 1 min to allow it to adapt to its new environment. After that, the divider was withdrawn and behavior was video-recorded for 10 mins with high resolution. To determine “naïve” male (with no prior sexual experience) courtship behavior, male flies were kept isolated up until they were tested in an experimental chamber. To determine the “trained” male courtship behavior, each male was first individually placed into an experimental chamber together with a mated female for 1 hour and then isolated for 30 mins prior to recording. Keeping males isolated for 30 mins prior to testing allowed for the measurement of short-term memory (STM) performance (Kamyshev et al., 1999; McBride et al., 2005). The courtship index (CI) of naïve and trained experimental male flies was calculated as a percentage of time a male spent performing courtship behavior during the recorded 10 min. The memory index (MI) was calculated as: [100(1-(CI of trained male/mean of CI of naïve male))] (Kamyshev et al., 1999; Lim et al., 2018).

### Affinity-purified mass spectrometry

Protein lysates were prepared from approximately 100 adult fly heads of wild-type, CS, or *acer^null^* flies. The heads were crushed on ice using a pestle in 0.5% NP-40 lysis buffer (50 mM Tris, 150 mM NaCl, 10% glycerol, 1 mM EDTA, pH 7.5) supplemented with protease inhibitors (Roche). The resulting lysates were clarified by centrifugation and the supernatants were collected. Acer protein purification was performed by incubating the lysate supernatants with primary anti-Acer antibody for 1 hour, followed by protein G Sepharose beads (GE healthcare life science, Cat #17061801) for 2 hours. Subsequently, the beads were washed with 0.5% NP-40 lysis buffer and then with buffer lacking NP-40. Further downstream processing and analysis were carried out at the SPARC BioCentre, The Hospital for Sick Children. The samples were first subjected to reduction by resuspending them in 100 µl of 50 mM ammonium bicarbonate and 10 mM DTT, followed by incubation at 60°C for 1 hour. Alkylation was performed using 20 mM iodoacetamide at room temperature in the dark for 45 minutes. Next, trypsin digestion was carried out overnight at 37°C using 2 µg of Pierce trypsin per sample at a ratio of 1:50-1:100. The resulting peptides were lyophilized and desalted using Millipore C18 ZipTips. The peptides were then resuspended in 0.1% formic acid for analysis using a Thermo Scientific Q-Exactive HF-X mass spectrometer coupled with an EASY nLC liquid chromatography system. Protein identification and quantification were performed using Proteome Discoverer version 2.2.0.388 and Scaffold 5.0.1 software, with a search against the Uniprot-*Drosophila*_*Melanogaste*r_Oct222018.fasta database and specific search parameters and modifications applied. SAINTexpress analysis was applied to the data to identify probable protein-protein interactions (Teo et al., 2014).

### Statistics

Statistical analyses were performed using GraphPad Prism software. For plaque staining and ELISA experiments, differences between two groups were analyzed using a two-tailed Student’s t-test. The Mann-Whitney *U* test was used to assess differences between groups in the TUNEL experiment and CI measurements obtained from the courtship conditioning assay experiment. To compare the MI between groups in the courtship conditioning assay experiment, the Kruskal-Wallis *H* test followed by Dunn’s multiple comparisons were utilized. The significance level was set at α < 0.05 for all tests. For gene and protein expression analyses, differences between two groups were analyzed using a two-tailed Student’s t-test.

## Results

### *acer^PD^*mutant allele reduces brain cell death and rescues STM in Aβ42 expressing flies similar to an *acer^null^* allele

We previously demonstrated that captopril suppressed AD-related phenotypes in fly models of AD, whereby we used the GAL4-UAS system to drive expression of human AD-related transgenes, including *UAS-Aβ42* (Aβ42) in the fly CNS, using the pan-neuronal *elav*^C155^-Gal4 (*elav*^C155^) driver (Lee et al., 2020). Furthermore, we showed that the benefits of captopril are mediated through Acer inhibition. However, it was unclear whether captopril achieved this by specifically inhibiting its catalytic activity. To investigate this, we created an enzymatically inactive variant of Acer, henceforth referred to as Acer^PD^, and examined its ability to reproduce the beneficial outcomes observed with an *acer^null^* allele in Aβ42 expressing flies as previously characterized (Lee et al., 2020). The coordination of zinc ions to histidine residues within the metalloprotease consensus motif, HExxH, present in mammalian ACE is essential for its catalytic function, and mutation of these histidine residues to lysine is known to disrupt ACE enzymatic activity (Jaspard & Alhenc-Gelas, 1995; Williams et al., 1994). We predicted that similar mutations in the Acer HExxH motif would disrupt its catalytic activity. Therefore, we aimed to introduce specific genomic single-nucleotide polymorphisms (SNPs) in the *acer* gene that result in the conversion of histidine residues to lysine within the HExxH motif. To introduce these mutations, we utilized a modified technique known as scarless gene editing that combines CRISPR-Cas9 mediated HDR with a visible marker to facilitate screening for successful HDR events. As illustrated in Figure 1A, a DsRed marker cassette was incorporated into the fly genome, along with the intended mutations H425K and H429K by HDR using a donor template. The cassette was designed to be expressed in fly eyes and is flanked by PBac transposon ends so that after successful HDR, a single cross to PBac transposase flies removed the cassette leaving behind only the mutations of interest. Identified *acer* mutants were sequenced to confirm gene editing and precise excision of the DsRed cassette and western blot analysis was performed to measure Acer^PD^ protein levels and compared to Acer^WT^ to ensure that the mutations did not alter protein expression levels (Fig. 1B).

**Figure 1.**
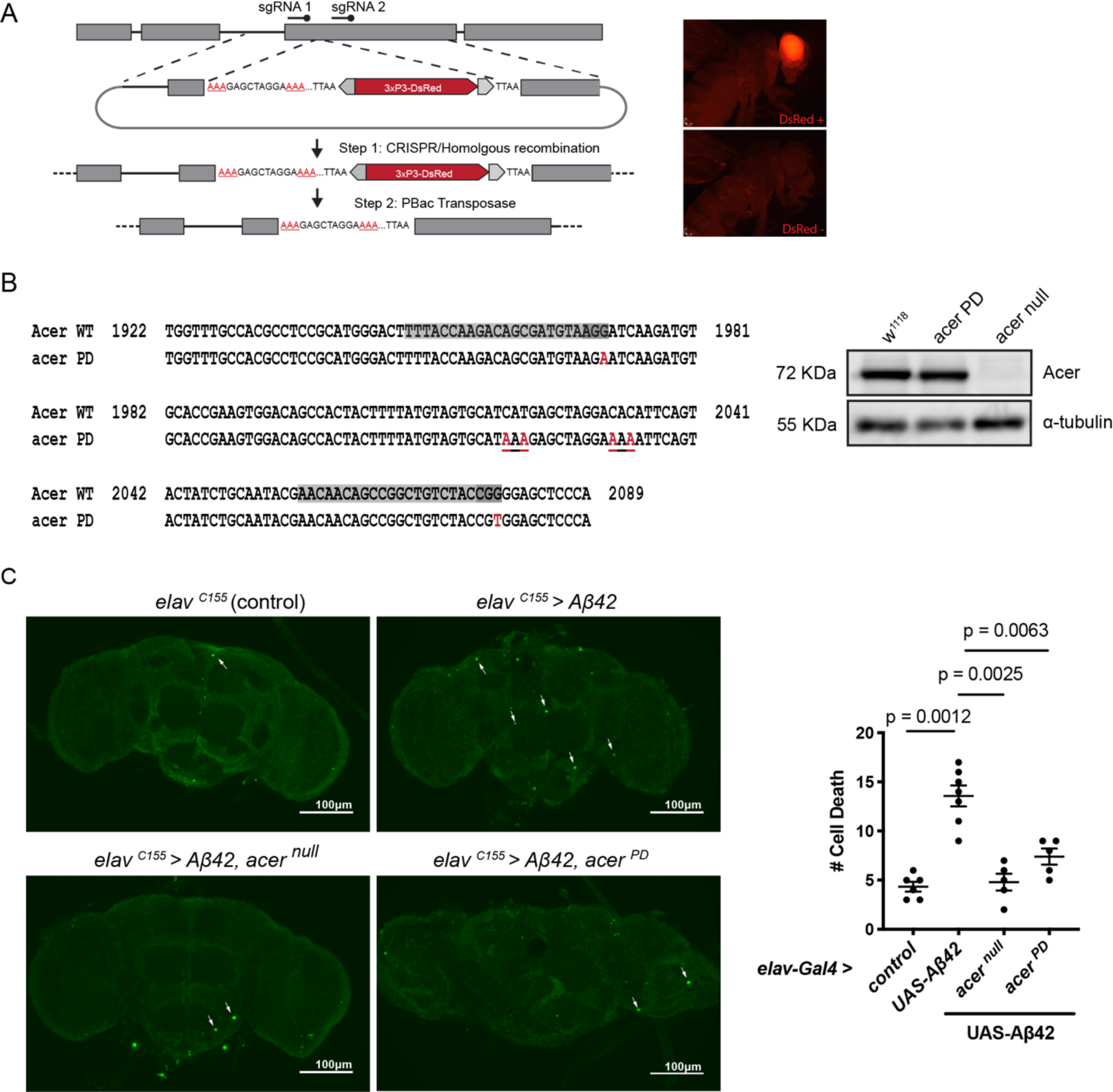
An *acer^PD^* mutation reduced brain cell death in Aβ42 flies similar to an *acer^null^* mutation. (**A**) Scarless gene editing scheme followed to introduce H375K and H379K mutations in *acer*. Mutations introduced by CRISPR-Cas9 HDR, including DsRed cassette. Following successful HDR, screened flies were crossed to PBac transposase flies resulting in precise excision of the DsRed cassette. Successful HDR events were screened by selecting flies positive for DsRed expression in fly eyes. Images represent fluorescence images of flies positive or negative for DsRed. (**B**) Sequence alignment of *Acer* and *acer^PD^*. Highlighted in light and dark gray are the sgRNA sequences and PAM sites, respectively. Underlined sequences are H375K and H379K mutations with point mutations highlighted in red. Intended silent mutations introduced in PAM sites are represented in blue. WB analysis of Acer protein levels in *w^1118^* control, *acer^PD^* and *acer^null^* flies; ɑ-tubulin used as a loading control. (**C**) Representative confocal images of TUNEL labeling in dissected brains from 4-week-old control flies, Aβ42 flies and Aβ42 flies carrying an *acer^null^* or *acer^PD^* mutation; arrows point to TUNEL positive cells (labeled in green), and quantification of TUNEL positive cells per imaged brain in the different genetic backgrounds. Data represent quantification of total cell death per brain; bars represent means ± SEM.

The *acer^PD^* mutant allele was then recombined with the *Aβ42* transgene to study the effects of the mutation on the pathological effects associated with Aβ42 expression in aged 4-week-old flies. We first examined brain cell death using TUNEL analysis. In line with previous findings (Finelli et al., 2004; Iijima et al., 2004; Lee et al., 2020), we show that expression of Aβ42 resulted in a significant increase in cell death within the adult brain compared to control (*elav* alone) flies. We also show that the presence of an *acer* null mutation in Aβ42 expressing flies significantly reduced the level of cell death as previously shown (Lee et al., 2020) (Fig. 1C). Similarly, we found that the *acer^PD^* mutant allele in Aβ42 expressing flies led to a significant reduction in the level of cell death, similar to that in control (*elav* alone) flies (Fig. 1C).

We next looked to see whether the *acer^PD^* mutant allele can rescue STM defects observed in Aβ42 expressing flies as we previously found with the *acer^null^* allele (Lee et al., 2020). We examined memory performance using a widely used conditioned courtship suppression assay (see section “Materials and Methods” for details) (Fig. 2A) (Griffith & Ejima, 2009; Kamyshev et al., 1999; Siegel & Hall, 1979). Courtship indices (upper graph) and corresponding memory indices (lower graph) for evaluated genotypes are presented in Fig. 2B. As expected, trained control (*elav* alone) males showed a significant reduction in courtship activity compared to their naïve counterparts, whereas trained Aβ42 expressing males did not (Fig. 2B, upper graph), which reflects a significant difference between their MI (Fig. 2B, lower graph). In turn, the presence of an *acer^null^* allele in Aβ42 expressing flies led to a significant reduction in male courtship activity of trained males compared to their naïve counterparts (Fig. 2B, upper graph) and resulted in a rescue of STM (Fig. 2B, lower graph). Interestingly, a similar finding was found with the *acer^PD^* mutant allele whereby trained Aβ42 expressing males homozygous for the *acer^PD^* allele displayed a significant reduction in their courtship after training (Fig. 2B, upper graph) and their MI was comparable to that observed in control (*elav^C155^* alone) flies (Fig. 2B, lower graph). Together, these data suggest that inhibiting the catalytic activity of Acer is necessary and sufficient to rescue AD-related STM deficit in Aβ42 expressing flies. Furthermore, it indicates that the processing of downstream target(s) of Acer is involved in mediating AD-related pathology.

**Figure 2.**
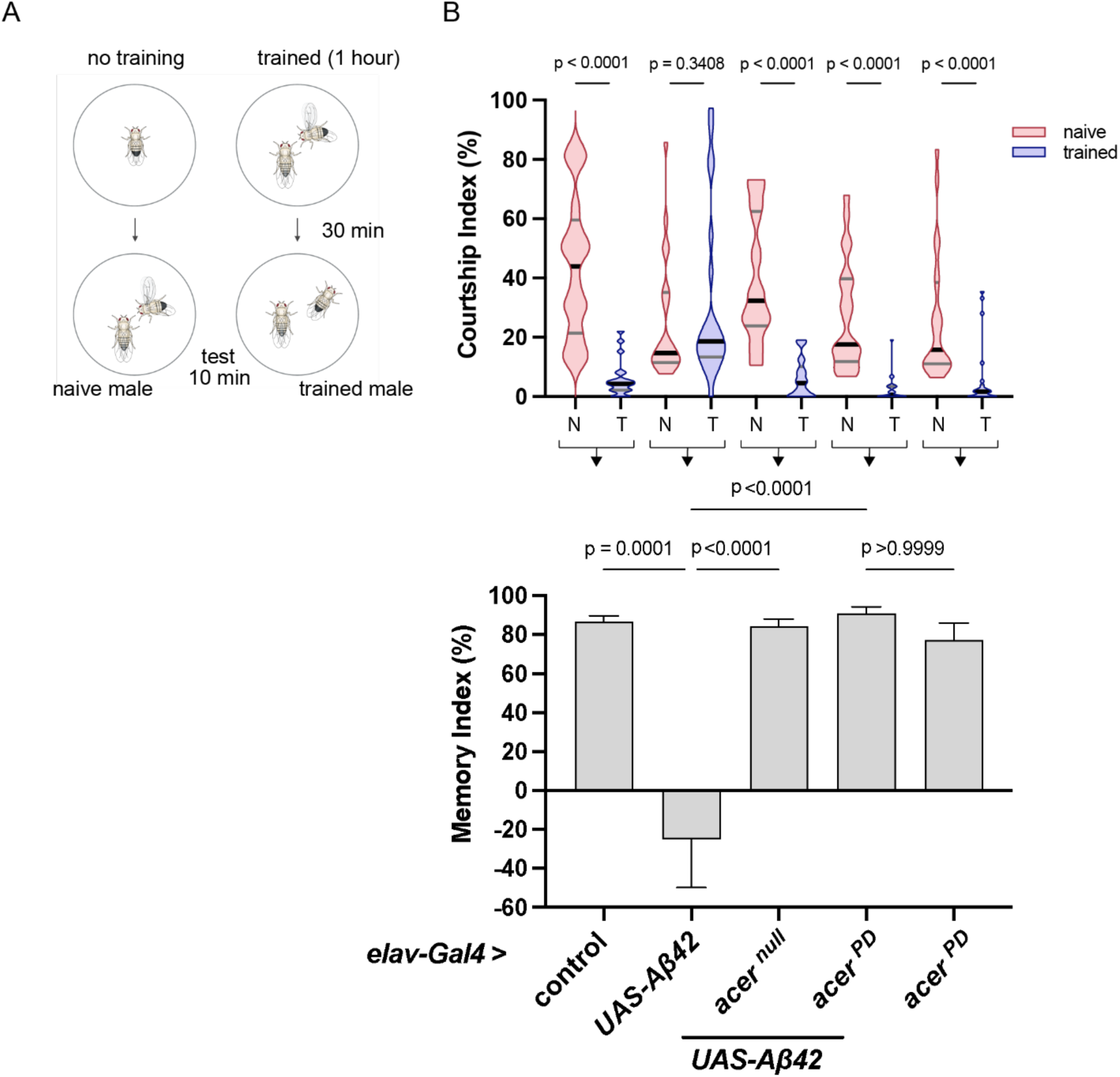
An *acer^PD^* mutation rescues STM in Aβ42 expressing flies similar to an *acer^null^*mutation. (**A**) Illustration of courtship conditioning assay. Two part – naïve male courtship behavior recorded for 10 min with no prior training; trained male is exposed to mated female 1 hour during training phase, then separated for 30 mins before testing and recording courtship behavior toward another mated female. (**B**) Upper graph: courtship indexes (CIs) for naïve (N) and trained (T) 4-week-old control (*elav^C155^* alone) flies, Aβ42 expressing flies, Aβ42 expressing flies homozygous for *acer^null^*or *acer^PD^* allele and *acer^PD^* allele in control background. N and T male CIs were compared within each genotype using two-tailed Mann-Whitney *U* tests. Data presented as a truncated violin plot of frequency distribution with lines: median (black), first and third quartiles (grey). Lower graph: memory indexes (MIs) calculated from respective CIs of N and T males (bracketed arrow) for corresponding genotypes. Memory between genotypes was compared using Kruskal-Wallis tests (H = 40.64, *p* < 0.0001) followed by Dunn’s multiple comparisons, with data presented as mean ± SEM; n > 21 per genotype.

### Acer interacts with CG2233 and regulates its protein levels

To identify Acer targets, we applied an unbiased approach by performing affinity purification mass-spectrometry (AP-MS). Protein lysates derived from adult heads of WT, *CS*flies were used to purify Acer using a specific antibody (Carhan et al., 2011), while lysates from *acer^null^*flies served as a control to account for non-specific interactors. Additionally, samples prepared from flies fed with captopril, an inhibitor of Acer (Houard et al., 1998), were included to help narrow down potential hits, as it allowed us to identify protein interactors that were hindered by captopril binding in a competitive manner. Therefore, any identified interactors that presented lower abundance levels in WT+captopril samples compared to WT samples suggested that these proteins interacted with the catalytic domain of Acer and that captopril had interfered with their interaction (Fig. 3). Proteins identified that fall under this criterion include CG2233, Photoreceptor dehydrogenase (Pdh) and ATPsynbeta. CG2233, according to FlyBase is an uncharacterized gene/protein, Pdh encodes a retinal pigment cell dehydrogenase involved in retinol metabolism, and ATPsynbeta is a mitochondrial membrane ATP synthase.

**Figure 3.**
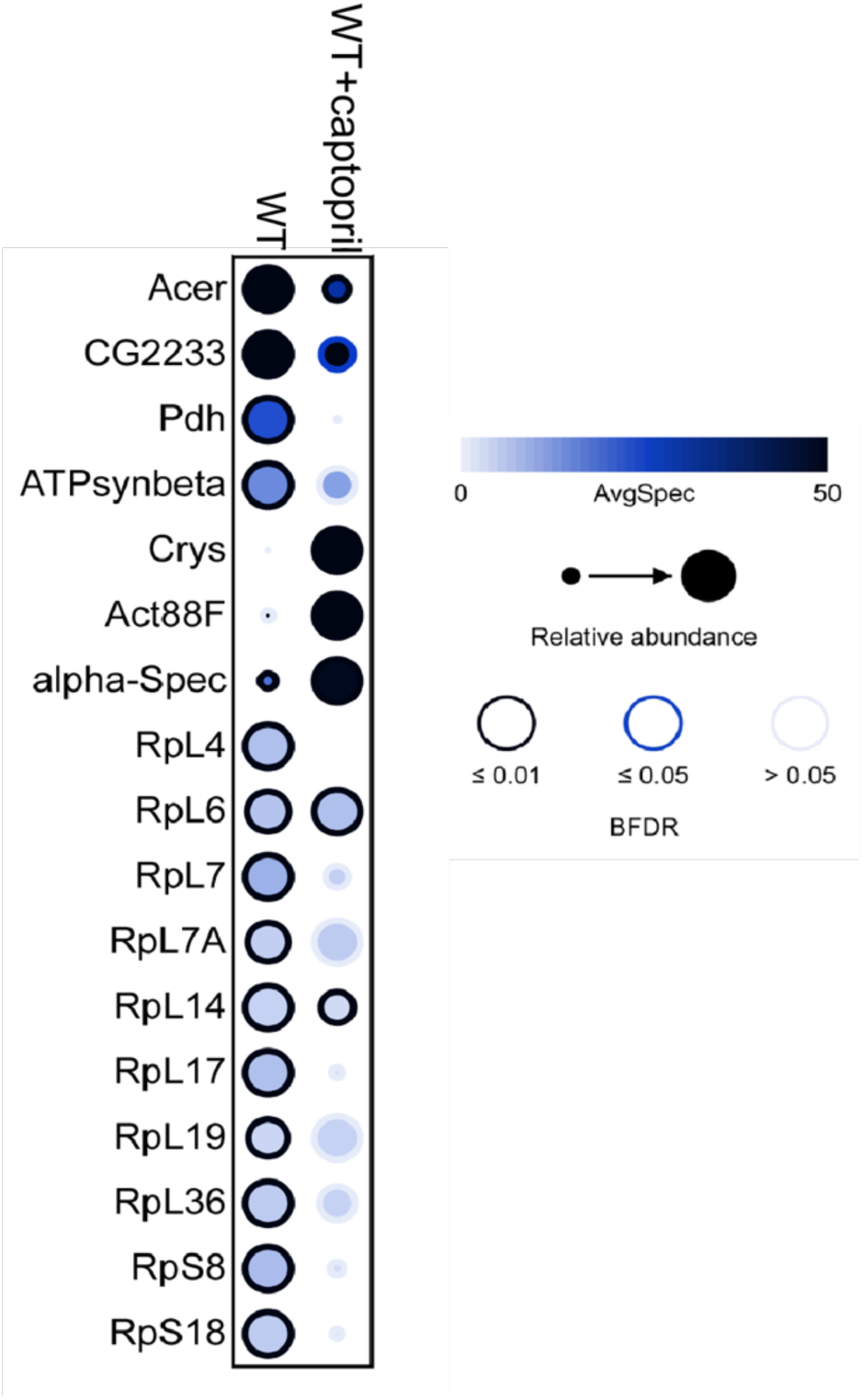
Acer protein interactors indentified through AP-MS. Top interacting proteins of Acer from AP-MS determined using the spectral count of WT and WT+Captopril samples normalized to Acer null samples. SAINTexpress analysis reports Bayesian false discovery rate (BFDR) used to determine significance. Data displayed by dot plot graph representing average spectral count, relative abundance and BFDR value.

Notably, while all three proteins were found to have a high SAINT score, only CG2233 was consistently identified as a strong potential interactor in two independent AP-MS experiments (data not shown). Furthermore, the addition of captopril appeared to diminish the interaction between CG2233 and Acer (Fig. 3). As such, CG2233 was considered a promising initial candidate for further investigation leading us to explore a functional interaction between CG2233 and Acer. We aimed to determine: (1) whether Acer plays a role in regulating CG2233 protein levels, (2) whether the levels are altered in the Aβ42 background compared to control, and (3) whether Acer influenced this change. To achieve these aims, we first generated a polyclonal antibody against CG2233 purified peptide and used it to measure CG2233 in different genetic backgrounds including control (*elav* alone) flies carrying *acer^WT^, acer^null^* or *acer^PD^*allele and Aβ42 expressing flies carrying *acer^WT^, acer^null^*or *acer^PD^* allele. (Fig. 4). We found that the level of CG2233 was significantly decreased in Aβ42 expressing flies when compared to control flies (*elav* alone). Furthermore, CG2233 levels were significantly increased in *acer^null^* and *acer^PD^* mutant flies in the control background (Fig. 4). However, this increase in fold change was no longer observed in flies carrying either the *acer^null^* or *acer^PD^* allele in the presence of Aβ42 expression. Nonetheless, these results suggest that Acer plays a role in regulating CG2233 levels and that inhibition of Acer results in an increase of CG2233. Additionally, these results also suggest that the levels of CG2233 are influenced by Aβ42 expression.

**Figure 4.**
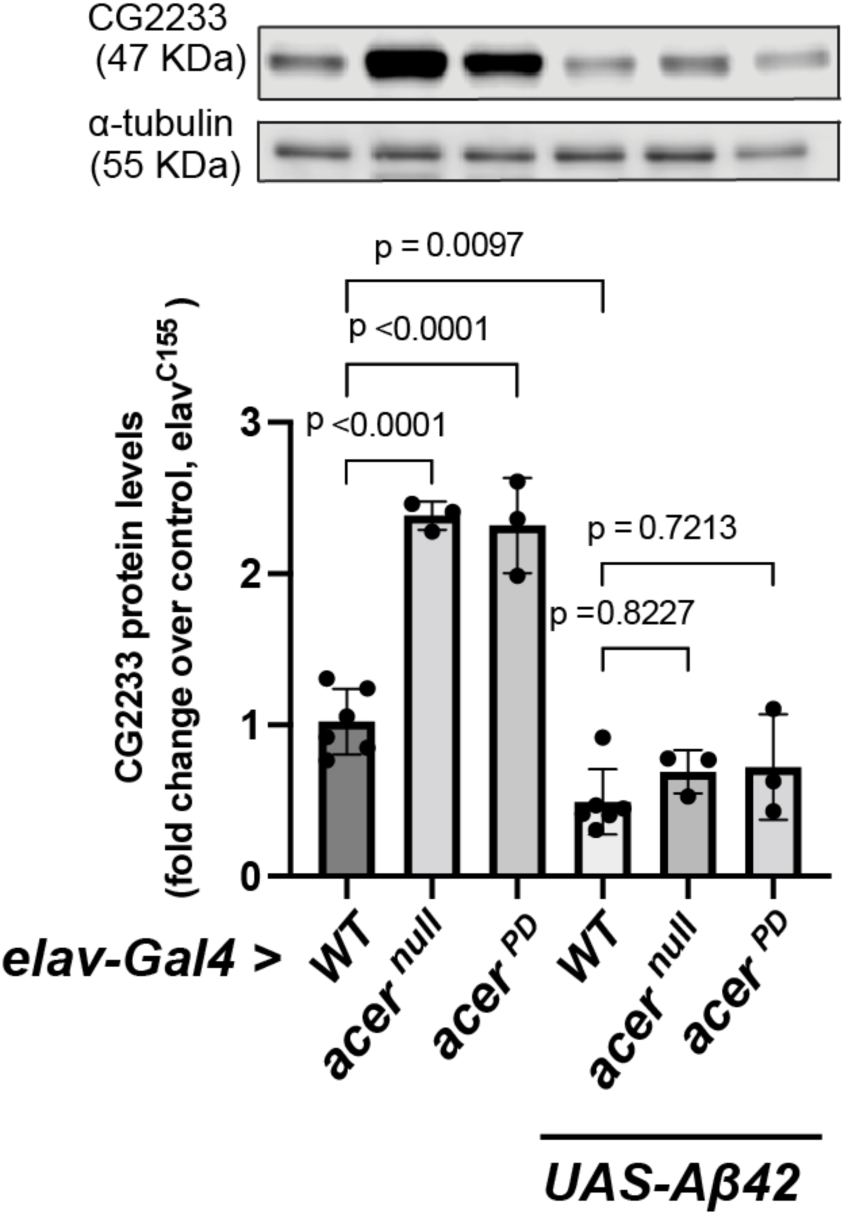
Acer regulates CG2233 protein levels. Representative WB and quantification of CG2233 protein levels measured from adult heads of 4-week-old control (*elav^C155^* alone) flies, control flies homozygous for *acer^null^ or acer^PD^* allele, Aβ42 expressing flies and Aβ42 expressing flies homozygous for *acer^null^ or acer^PD^* allele. CG2233 levels normalized to ɑ-tubulin in each genotype and presented as fold change over control. Fold change across different genotypes were compared using one-way ANOVA followed by Tukey’s multiple comparisons test, and data presents mean ± SD, n ≥ 3.

To investigate a potential role for CG2233 in pathways underlying Aβ42 pathology, we generated a *CG2233* null mutation, *CG2233^Δ1353^*, and evaluated its effects on Aβ42 induced phenotypes. A null mutant was generated using CRISPR-Cas9 NHEJ, whereby two sgRNAs were designed to target and remove the majority of the *CG2233* open reading frame (ORF) (illustrated Fig. 5A), resulting in a 1353 base pair deletion that removed 76% of the ORF. WB analysis revealed no protein, indicating that the mutation functioned as null (Fig. 5B). Of note, these flies are homozygous viable and display no obvious phenotypes. We first evaluated the effects of the mutation on the level of cell death in 4-week-old Aβ42 expressing fly brains. Interestingly, we found that the *CG2233^Δ1353^* mutation resulted in an increase of the level of cell death observed in brains of Aβ42 expressing flies (Fig. 5C). In contrast, the mutation had no effect on the level of cell death in the control background (*elav^C155^*alone), suggesting that *CG2233^Δ1353^* specifically enhances Aβ42 induced cell death. We also evaluated the effect of the *CG2233^Δ1353^* mutation on the STM defect induced by Aβ42 expression. Of note, Aβ42 flies exhibited a complete memory loss at 4 weeks as determined by a one-sample Wilcoxon signed-rank test (*W* = 21.00, *p =* 0.750) (Fig. 2B lower graph),such that we would not be able to measure any effects due to the presence of the *CG2233^Δ1353^* mutation. Therefore, we decided to perform this assay at 3 weeks of age when -week-old Aβ42 flies might display a less severe phenotype, allowing us to determine if loss of CG2233 could enhance this phenotype. However, we found that even at 3 weeks, Aβ42 expressing flies also displayed complete STM loss; a one-sample Wilcoxon signed-rank test revealed no significance (*W* = 113.0, *p* = 0.1336). Interestingly, however, we found that the *CG2233^Δ1353^* mutation resulted in a complete rescue of the STM defect exhibited in Aβ42 expressing flies. Aβ42 expressing flies carrying the *CG2233^Δ1353^*mutation show a significant reduction in their CI after training compared to the CI of respective untrained naïve male flies (Fig.6A upper graph), and their MI was comparable to the MI of control (*elav^C155^*alone) flies (Fig.6A lower graph). This ability of CG2233 to rescue the memory defect was surprising given that loss of CG2233 enhanced the levels of cell death observed in 4-week-old Aβ42 expressing flies.. Therefore, we ext asked if the ability of CG2233 mutants to rescue the memory defects in AB42 expressing flies persisted in older flies. Our results show that the *CG2233^Δ1353^* mutation failed to rescue STM loss in 4-week-old Aβ42 flies (Fig.5B lower graph). Aβ42 expressing male flies carrying the *CG2233^Δ1353^* mutation did not exhibit a reduction in their CI after training compared to untrained naïve males (Fig.5B upper graph), and their calculated MI has not significantly different from Aβ42 control flies (Fig.6B lower graph).

**Figure 5.**
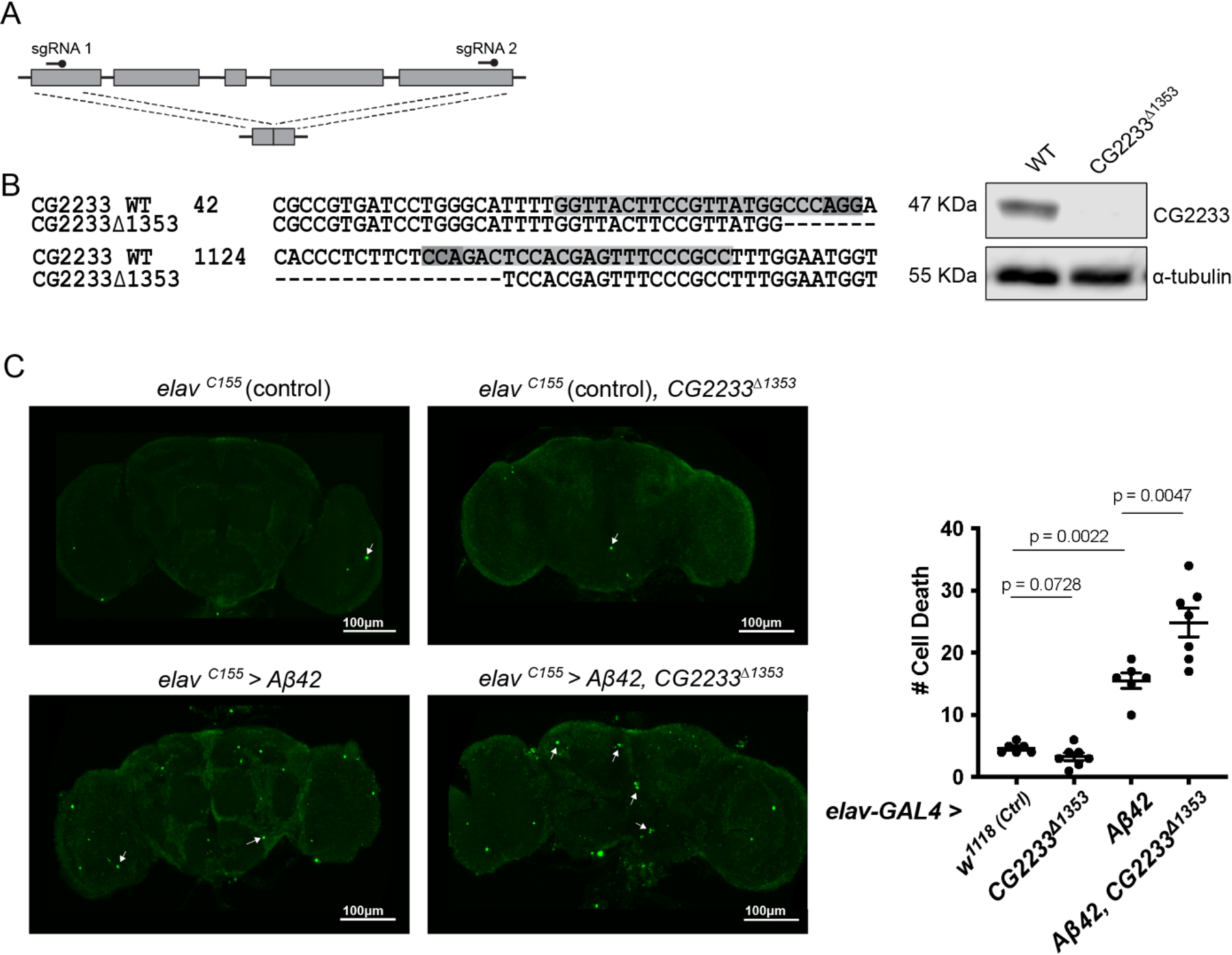
*CG2233^Δ1353^* mutation enhances brain cell death in Aβ42 expressing flies. (A) Schematic representation of two sgRNA designed to cleave at exon 1 and 5. (B) Sequence of mutants highlighting gRNA1 and 2 sequence (grey), PAM sites (dark grey) and deletion region, and WB analysis of CG2233 in WT and mutant lines with polyclonal antibody against CG2233; ɑ-tubulin used as a loading control. (B) Representative confocal images of TUNEL labeling in dissected brains from 4-week-old control (*elav^C155^* alone) flies, Aβ42 expressing flies, and Aβ42 expressing flies carrying the *CG2233^Δ1353^* mutation; arrows point to TUNEL positive cells (labeled in green), and quantification of TUNEL positive cells per imaged brain in the different genetic backgrounds. Data represent the quantification of total cell death per brain; data represents means ± SEM, n > 6.

**Figure 6.**
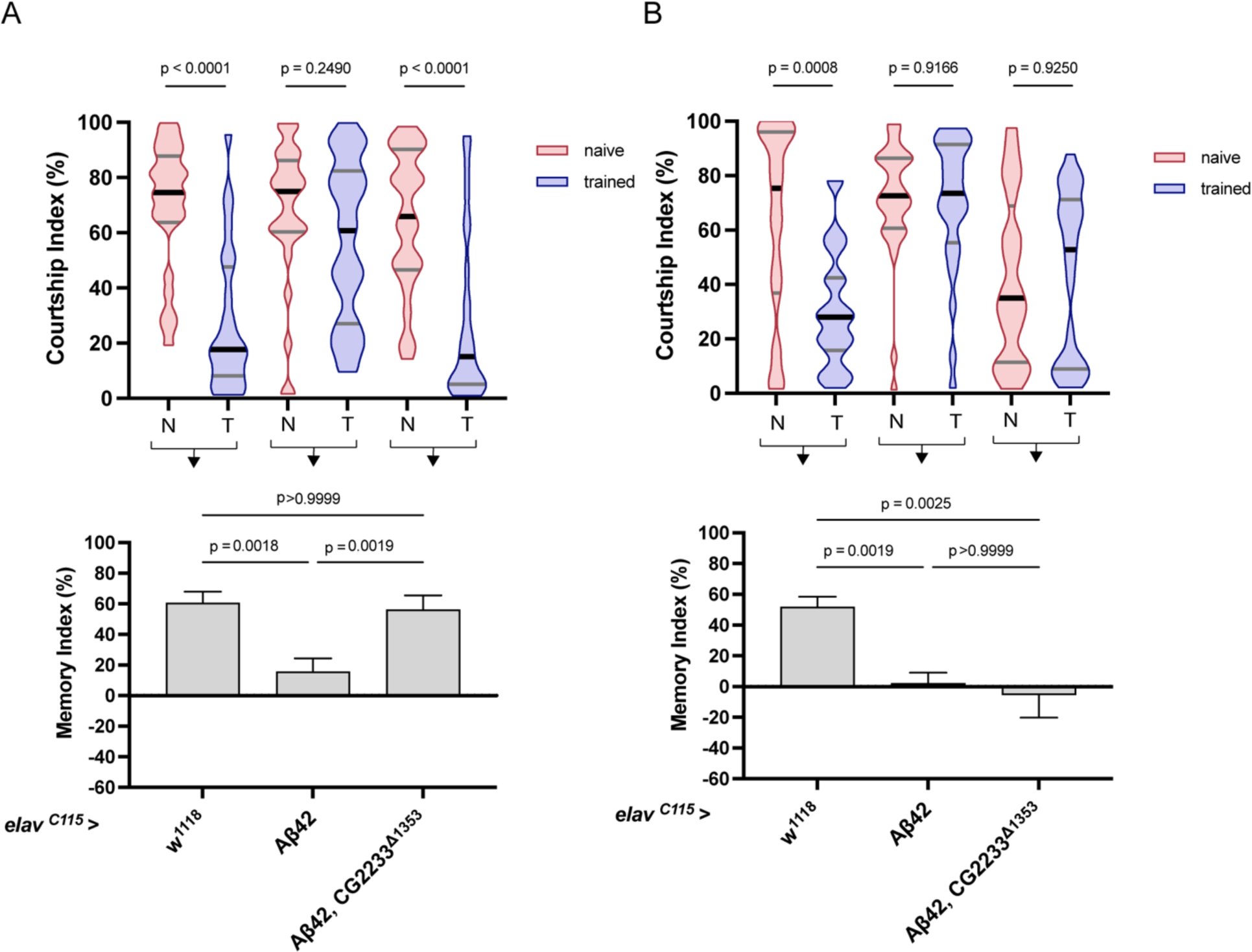
CG2233^Δ1353^ mutation rescues STM in Aβ42 expressing flies at 3 weeks but not 4 weeks and instead disrupts normal courtship behavior. **(A)** Upper graph: Courtship indexes (CIs) for naïve (N) and trained (T) 3-week-old control (*elav^C155^* alone) flies, Aβ42 expressing flies, and Aβ42 expressing flies carrying the *CG2233^Δ1353^* mutation. N and T male CIs were compared within each genotype using two-tailed Mann-Whitney *U* tests. Data presented as a truncated violin plot of frequency distribution with lines: median (black), first and third quartiles (grey). Lower graph: memory indexes (MIs) calculated from respective CIs of N and T males (bracketed arrow) for corresponding genotypes. Memory between genotypes was compared using Kruskal-Wallis tests (H = 15.51, *p* = 0.0004) followed by Dunn’s multiple comparisons. Data presented as mean ± SEM; n > 25 per genotype. **(B)** Upper graph: CIs measured in 4-week-old male flies, lower graph: Mis for corresponding genotypes. Memory between genotypes was compared using Kruskal-Wallis tests (H = 15.39, *p* = 0.0005) followed by Dunn’s multiple comparisons, with data presented as mean ± SEM; n > 24 per genotype.

Notably, when we compared the CI of naïve 4-week-old males across the different genotypes (Fig.6B lower graph), we found a significant decrease in courtship activity in Aβ42 expressing male flies carrying the *CG2233^Δ1353^* mutation. A Mann Whitney two-tailed statistical test used revealed a significant difference between Aβ42 expressing flies carrying the *CG2233^Δ1353^* mutation and control (*U* = 209, *p* = 0.0191) and to Aβ42 expressing flies (*U* = 172.5, *p* = 0.0003). No significant difference was found between the CIs of control (*elav^C155^* alone) and Aβ42 expressing flies (*U* = 445, *p* = 0.9677). This suggests that the *CG2233^Δ1353^* mutation resulted in a disruption of normal male courtship behavior in Aβ42 expressing flies aged to 4 weeks. Previous research has reported a disruption in normal male courtship behavior as a consequence of neurodegeneration (Hu et al., 2014)., This finding may reflect an exacerbation of Aβ42 pathology due to loss of *CG2233* in the background of Aβ42 expressing flies. Therefore, while the CG2233 mutation failed to rescue STM in 4-week-old Aβ42 expressing flies it did result in a significant reduction in courtship behavior.

## Discussion

Although growing evidence supports the potential beneficial use of ACE-Is in AD, their mechanism of action in the context of the disease remains poorly understood. One challenging factor contributing to the knowledge gap is the inability to distinguish between their impact on blood pressure regulation and their direct influence on local RAS within the brain. Consequently, *Drosophila*, a model with an open circulatory system that also lacks a conserved RAS pathway (Fournier et al., 2012; Salzet et al., 2001), provides a unique opportunity to explore the link between ACE-Is and AD. Accordingly, we have been interested in evaluating the effects of ACE-Is and determining their mechanism of action using *Drosophila* that express human AD-related transgenes. We have previously shown that captopril, an ACE-I, reduced levels of brain cell death and rescued memory deficits in these flies (Lee et al., 2020). Furthermore, we demonstrated that a null mutation in *acer,* a *Drosophila* homolog of *Ace*, is sufficient to recapitulate these effects, suggesting that Acer is the target of captopril. We also showed that the beneficial effects are independent of changes in Aβ42 levels or plaque load (Lee et al., 2020). Of note, while ACE homologs have been identified in *Drosophila*, other components of the RAS are not conserved, suggesting a potentially novel function of ACE linked to AD. To unravel this potential function, we characterized the function of Acer in Aβ42 expressing flies. We showed that a predicted inactive form of Acer was sufficient to reduce cell death in the brains of Aβ42 expressing flies and rescue observed STM loss, therefore establishing that the enzymatic activity of Acer is important in mediating Aβ42-related phenotypes. Nonetheless, the function of Acer remains poorly studied, limiting our understanding of how its inhibition would suppress AD-related phenotypes. Previous studies aimed at elucidating its function either through ACE-Is or genetic nulls have provided some insights. A study by Carhan et al. (2011) revealed a role for Acer in sleep regulation, which had been suggested by the cyclical expression of *Acer* in adult heads regulated by the circadian gene (*clock*) (Carhan et al., 2011; McDonald & Rosbash, 2001). Although the precise mechanisms underlying the role of Acer in sleep regulation are not yet fully understood, it has been hypothesized to be associated with alterations in metabolic processes (Glover et al., 2019). Acer is prominently expressed in the fat body, which has been shown to regulate metabolism, hormone secretion, and complex behaviors such as sleep (Arrese & Soulages, 2010; Yurgel et al., 2018). Indeed, a recent study by Glover et al. (2019) determined a potential role for Acer in metabolism, particularly in glycogen storage. This opens up an intriguing avenue for further research to study the link between Acer and AD-related phenotypes. AD is notably characterized by a decline in glucose metabolism that is believed to contribute to disease pathogenesis (Kumar et al., 2022). A decrease in metabolism is attributed to reduced uptake of glucose by the brain. Furthermore, Niccoli et al. (2016) showed that increasing glucose uptake in neurons mitigated neurodegeneration and extended lifespan in a *Drosophila* AD model. Consequently, it warrants investigation to determine whether Acer may play a role in sustaining proper glucose metabolism within the brains of these flies.

As an alternative approach to identify novel roles for Acer we took an unbiased approach to identify potential protein interactors. We identified and selected CG2233 to be a promising prospective target to investigate. Subsequently, CG2233 protein levels were investigated under various conditions and were found to be regulated by Acer and Aβ42 expression, respectively; inhibition of Acer results in a significant increase in CG2233 protein levels, and Aβ42 expression results in significant downregulation of CG2233 protein level compared to control background. These findings provided compelling evidence for a functional relationship between Acer and CG2233 and suggested the involvement of CG2233 in Aβ42-related phenotypes. Consequently, a role for CG2233 was examined in Aβ42 expressing flies by introducing a *CG2233* null mutation in the background of Aβ42 expression. Evaluating flies at 4 weeks revealed that the mutation enhanced brain cell death and disrupted normal courtship behavior, while at 3 weeks, it rescued STM defects, suggesting that CG2233 possesses a dual mode of action that is time-dependent. At early stages in Aβ42 pathology, the knock-out of CG2233 appeared to be beneficial, however, at later stages, it appeared to be detrimental. Taken together, data collected has revealed an important link between Acer-CG2233 and Aβ42 pathology. Building upon this finding, it is possible to examine what is known of CG2233 to elucidate underlying cellular pathways modulated by the Acer-CG2233 interaction. While a direct homolog of CG2233 may not exist in mammals, downstream pathways shared by CG2233 and Acer may be conserved and could provide insightful knowledge on the function of mammalian ACE and AD. For instance, links were found between CG2233 and innate immune responses (Troha et al., 2018; Troha et al., 2019; Ullastres et al., 2021). In a study aimed at identifying novel genomic regulatory regions occupied by transposable elements that are responsible for interindividual differences in response to immune challenges in *Drosophila*, several regions, including those that regulate *CG2233* expression, were identified (Ullastres et al., 2021). This same study showed that the knockdown of *CG2233* results in increased survival rates after bacterial infection. This link becomes increasingly noteworthy when considered alongside previous research on ACE-Is in mouse models of AD. Studies demonstrated that ACE-Is reduced neuroinflammation, a critical pathological factor of AD (Dong et al., 2011; AbdAlla et al., 2013; Asraf et al., 2018; Torika et al., 2016). While the effects of ACE-Is are attributed to RAS inhibition, exploring the potential connection between Acer and the innate immune system via CG2233 regulation could possibly reveal a role for ACE in AD unrelated to RAS regulation. Collectively, this study offers valuable insights into the mechanism of ACE inhibition in a *Drosophila* model of AD. It also presents a promising avenue for further investigation through the characterization of CG2233, shedding light on potential pathways associated with AD pathology that implicate ACE.

## Acknowledgements

This work was supported by a grant from the Canadian Institutes of Health Research (PJT153063to G.L.B.

